# AGO HITS-CLIP in Adipose Tissue Reveals miR-29 as a Post-Transcriptional Regulator of Leptin

**DOI:** 10.1101/2020.03.13.991422

**Authors:** Sean O’Connor, Elisabeth Murphy, Sarah K. Szwed, Matt Kanke, François Marchildon, Robert B. Darnell, Praveen Sethupathy, Paul Cohen

**Author notes:** Corresponding Author Paul Cohen, M.D., Ph.D., The Rockefeller University, Laboratory of Molecular Metabolism, 1230 York Ave. New York, NY 10065.

## Abstract

MicroRNAs (miRNAs) are short, non-coding RNAs that associate with Argonaute (AGO) to regulate mRNA stability and translation. While individual miRNAs have been shown to play important roles in white and brown adipose tissue in normal physiology and disease^1,2,3^, a comprehensive analysis of miRNA activity in these tissues has not been performed. We used high-throughput sequencing of RNA isolated by crosslinking immunoprecipitation (HITS-CLIP) to comprehensively characterize the network of high-confidence, *in vivo* mRNA:miRNA interactions across white and brown fat, revealing over 100,000 unique miRNA binding sites. Targets for each miRNA were ranked to generate a catalog of miRNA binding activity, and the miR-29 family emerged as a top regulator of adipose tissue gene expression. Among the top targets of miR-29 was leptin, an adipocyte-derived hormone that acts on the brain to regulate food intake and energy expenditure^4^. Two independent miR-29 binding sites in the leptin 3’-UTR were validated using luciferase assays, and miR-29 gain and loss-of-function modulated leptin mRNA and protein secretion in primary adipocytes. In mice, miR-29 abundance inversely correlated with leptin levels in two independent models of obesity. This work represents the only experimentally generated miRNA targetome in adipose tissue and identifies the first known post-transcriptional regulator of leptin. Future work aimed at manipulating miR-29:leptin binding may provide a therapeutic opportunity to treat obesity and its sequelae.

---

Much of the work on adipocyte development and function over the last several decades has focused on transcriptional regulators. These include PPARG2, which is necessary and sufficient for the development and maintenance of white and brown adipocytes^5^, and PRDM16, which regulates brown and beige adipocyte identity^6,7^. More recently, it has become clear that post-transcriptional regulators also play a key role in influencing adipocyte phenotype. Studies with adipose-specific dicer KO mice demonstrate that microRNAs (miRNAs) are essential in regulating adipogenesis, insulin sensitivity and non-shivering thermogenesis^8,9^. Several individual miRNAs have been shown to contribute to these phenotypic effects. miR-133 represses BAT function by directly targeting *Prdm16* and inhibiting expression of thermogenic genes^2^. miR-196a induces browning of WAT by targeting *Hoxc8*^1^, and miR-26 protects mice from diet-induced obesity by blocking adipogenesis^10^. These studies, and most others in adipose tissue, rely on computational predictions and low-throughput validation to identify miRNA targets. Although high-throughput crosslinking techniques to map the miRNA targetome have been applied to liver, brain, heart and other tissues^11,12,13^, thus far, no comprehensive targetome has been described for adipose tissue.

To profile the complete network of *in vivo*, adipose-specific miRNA binding activity across both brown and white fat, we used AGO HITS-CLIP on interscapular brown adipose tissue (iBAT) and epididymal white adipose tissue (eWAT). Following UV crosslinking, AGO:RNA complexes were immunoprecipitated (Supplementary Fig. 1a), radiolabeled, and separated by gel electrophoresis. iBAT and eWAT samples contained a range of RNA fragment lengths, illustrated by a dense smear on the autoradiograph, compared to IgG and non-crosslinked controls (Supplementary Fig. 1b). After excising AGO:RNA complexes of roughly 110 – 170 kDa and sequencing AGO-associated RNAs, iBAT and eWAT samples separated effectively by PCA (Fig. 1a) and showed low inter-sample variability by Pearson correlation (Supplementary Fig. 1c). mRNA tags (sequencing reads corresponding to AGO-associated mRNA fragments) were clustered into significant peaks revealing a total of 21,281 unique peaks across 6,717 genes. Approximately 53% of peaks localized to mRNA 3’-untranslated regions (UTRs), ∼40% localized to coding sequences (CDS) and the remaining 7-8% localized to 5’-UTRs, introns and other regions (Fig. 1b). Total unique peaks with greater than 2 tags were higher in iBAT than eWAT, although the proportion of peaks mapping to the 3’-UTR vs. CDS was roughly equivalent in both depots (Fig. 1c).

**Fig. 1.**
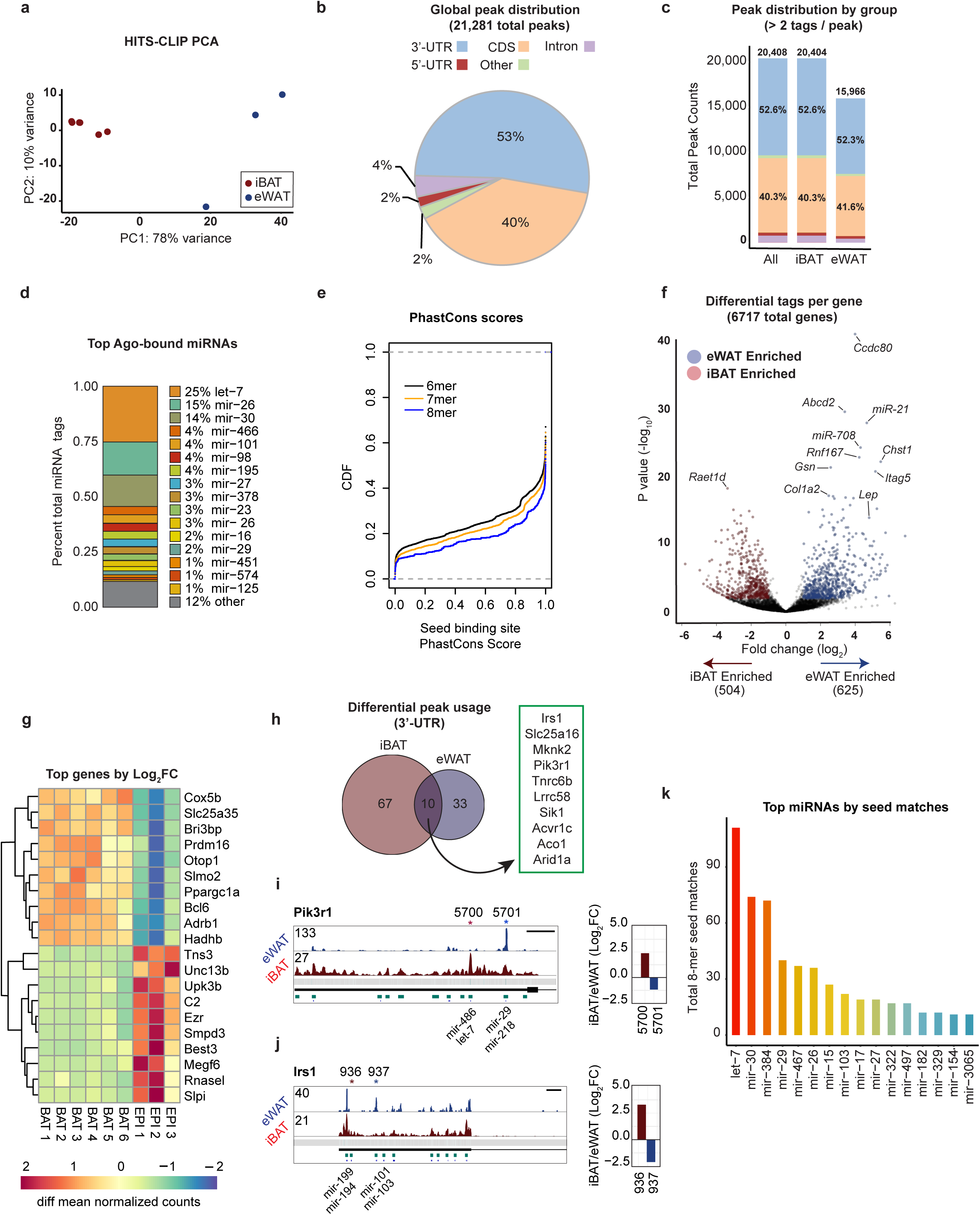
HITS-CLIP reveals miRNA bindings sites in iBAT and eWAT. **a**, PCA analysis of HITS-CLIP peaks from iBAT and eWAT (13-week old male mice, 2 mice pooled per group, n = 6 groups for BAT, n = 3 groups for eWAT). **b**, Global peak distribution by annotated region. **c**, Number of peaks with > 2 tags per peak, by annotated region, for iBAT and eWAT. **d**, Abundance of top 16 AGO-associated miRNAs as a percentage of total normalized miRNA tags. **e**, CDF plot of PhastCons scores from the top 1000 peaks (ranked by peak height), separated by seed match category. **f**, Volcano plot of iBAT vs. eWAT differential analysis (DESEQ2) of tags per gene. **g**, Heatmap of top 10 iBAT enriched and top 10 eWAT enriched genes by tag abundance. **h**, Venneuler plot: differential peak usage across depot types, subsetted to only include peaks in 3’UTRs. 77 genes contain peaks that are more highly bound by AGO in iBAT, 43 genes have peaks more highly bound in eWAT, and 10 genes contain both iBAT and eWAT enriched peaks. **i,j**, Gene track of the *Pik3r1* (**i**) and *Irs1* (**j**) loci showing tags plotted by genomic location. Turquoise bars below peaks represent half peak height of statistically significant peaks, and blue vertical bars represent miRNA binding sites. *significantly differential peak (vertically aligned), red = iBAT preferred, blue = eWAT preferred. Scale bar = 500 bp. Panels to the right of gene tracks show iBAT/eWAT (Log_2_ fold change) for mean tags in significantly differential peaks. **k**, Bar plot of most abundant 8-mer miRNA seed binding sites within the top 1000 peaks by peak height. Differential analysis of tags performed using DESEQ2. Significance cutoff: false discovery rate (FDR) < 0.05 and |Log_2_FC| > 1.

HITS-CLIP reads were next mapped to a library of mature miRNAs, resulting in 305 unique AGO-associated miRNAs (Extended Table 1). Among these, the 15 most abundant miRNAs comprised 88% of total miRNA tags in adipose tissue (Fig 1d). AGO-associated miRNAs were matched with binding sites by searching for seed sequence complementarity within peaks, then conservation across 26 species was assessed by PhastCons^14^ to assign a score to each miRNA seed sequence binding site and to 200 base pairs of DNA flanking each full miRNA binding site (Extended Table 2). Of a total of 138,187 miRNA:target interactions, 72% of seed binding sites were better conserved than their flanking regions. Cumulative distribution function (CDF) plots of PhastCons scores for miRNA seed sequence binding sites in the top 1000 significant peaks showed that, in aggregate, longer seed sequence matches are associated with increased conservation (Fig. 1e).

A differential analysis was performed on tags per gene (Fig. 1f) and tags per peak (Supplementary Fig. 1d,e) comparing eWAT and iBAT. *Lep* was among the mRNAs most bound by AGO in eWAT relative to iBAT (Fig 1f), and *Prdm16* and *Pgc1a* were among the top iBAT bound mRNAs (Fig. 1g). Next, to identify peaks with depot-specific regulation while accounting for differences in total AGO binding, peaks enriched in iBAT or eWAT were subsetted from genes with equal levels of total tags, resulting in 77 iBAT genes and 43 eWAT genes with depot-specific peaks (Fig. 1h). Of these, 10 genes contained both iBAT and eWAT enriched peaks, including *Pik3r1* (Fig. 1i) and *Irs1* (Fig. 1j), two genes involved in insulin signaling transduction. Interestingly, miR-218 is an eWAT enriched miRNA that binds to peak 5701 in *Pik3r1*, consistent with a potential role for miR-218 in promoting insulin resistance in visceral fat^15^.

To identify miRNAs that are most responsible for regulating gene expression in adipose tissue, miRNAs were ranked by high-confidence seed matches (8-mer binding sites within the top 1000 peaks). The top 2 ranked candidates, let-7 and miR-30, have been thoroughly studied in adipose tissue, both *in vivo* and *in vitro*^16,17,18,19^, and the third ranked candidate, miR-384 is not highly expressed (Extended Table 1). For this reason, we decided to focus on miR-29, the fourth-ranked candidate (Fig. 1k). miR-29 has been described previously as a regulator of the glucocorticoid receptor in adipocytes^20^, but has otherwise not been well studied in fat cells. Of the three miR-29 family members, miR-29a is 60-70 fold more abundant in adipose tissue than miR-29b and miR-29c (Supplementary Fig. 2a) and is expressed in both mature adipocytes and the stromal vascular fraction (SVF; Supplementary Fig. 2b). We performed a GO-term enrichment analysis of miR-29 targets with 8-mer seed matches and found that about half of the top pathways were related to ECM structure or insulin signaling, consistent with miR-29’s previously described roles in liver, cornea, muscle, and heart (Supplementary Fig. 2c)^21,22,23,24^. A closer examination of individual miR-29 targets, ranked by peak height, reveals that 4 of the top 10 genes have been previously validated in other cell types (Fig. 2a). To validate the 6 novel targets, miR-29a mimics were co-transfected into HEK-293A cells with a dual luciferase plasmid containing miRNA binding sites cloned into the 3’-UTR of firefly luciferase. For 5 of the novel binding sites, normalized luminescence was significantly decreased with miR-29a, but not a scrambled control (Fig. 2b-e,g). Mutations introduced within the miR-29 seed sequence binding sites were sufficient to block this effect. Among the top 10 ranked targets there was one 6-mer binding site, located in *Hadha*, which did not validate (Fig 2f).

**Fig. 2.**
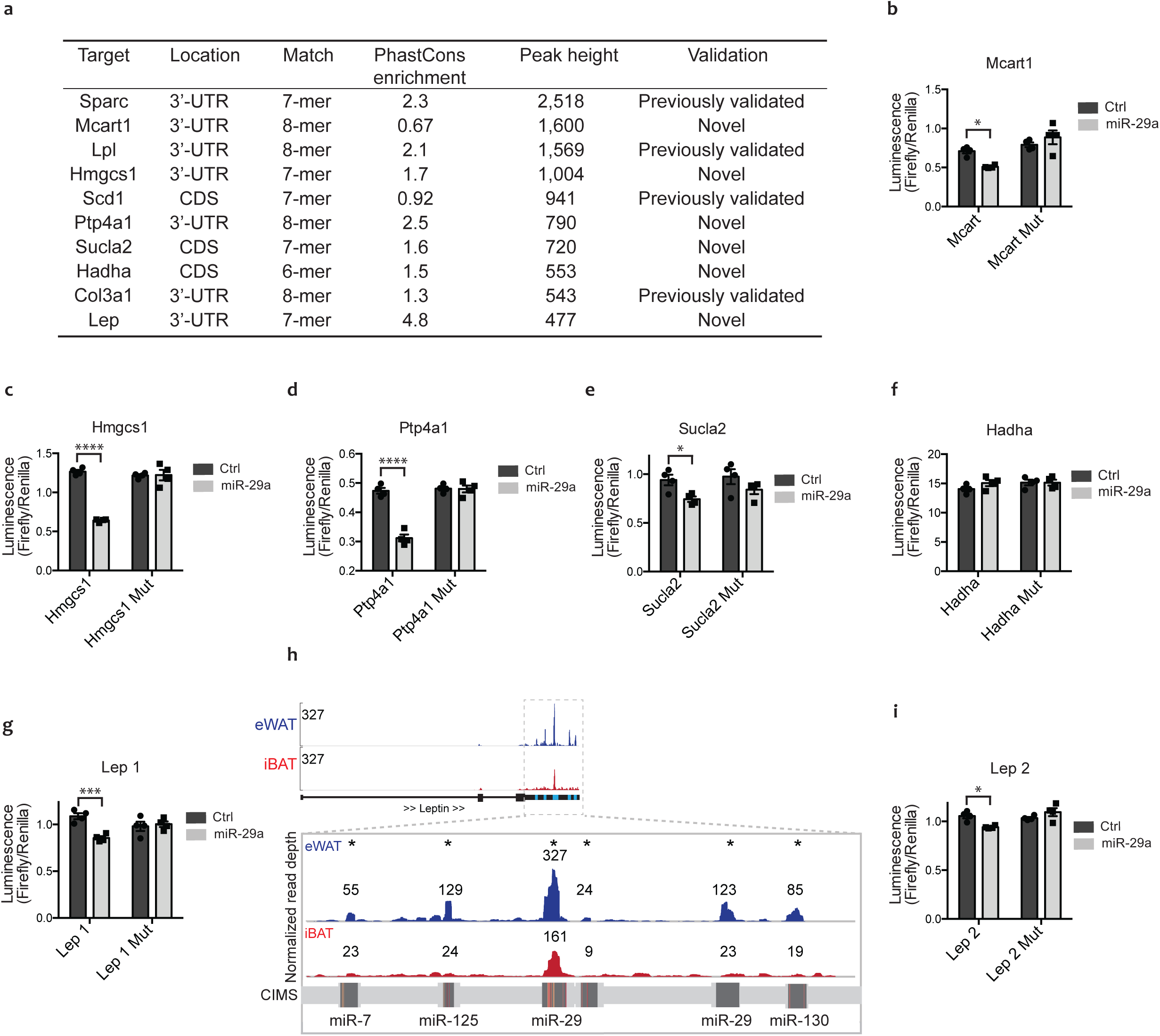
miR-29 has two binding sites in the leptin 3’-UTR. **a**, Table of top miR-29 targets, ranked by peak height, with previously validated binding sites noted^24,44,45,46^. **b-g,i**, Dual luciferase assay validation conducted by co-transfecting HEK-293A cells with plasmid containing luciferase enzymes and either a scrambled control miRNA or miR-29a. Luciferase activity is normalized to Renilla activity and presented as a ratio (n = 4). Mut = luciferase plasmid containing point mutations in seed sequence binding site. (**g**) and (**i**) refer to the miR-29 binding sites in leptin of peak heights 327 and 123, respectively. **h**, Gene track of the leptin locus showing tags plotted by genomic location relative to a maximum peak height of 327. Blue bars below peaks represent statistical significance. In insert, * represents statistically significant peaks and numbers indicate peak height. In the crosslinking induced mutation site (CIMS) track, red bars indicate robust mutation sites, and yellow bars indicate weak mutation sites. All results are presented as mean ±SEM. 2-way ANOVA used for miR-29a to ctrl comparisons (*p < 0.05, ***p < 0.0005, ****p < 0.00005)

Of miR-29’s novel targets, we focused on leptin, which plays a central role in energy homeostasis^4^. A map of the leptin locus shows that AGO HITS-CLIP tags fall almost exclusively within the 3’-UTR (Fig. 2h). Normalized tags cluster into 6 statistically significant peaks, 5 of which have complementarity with miRNA seeds. Many of the miRNA binding sites are marked by cross-linking induced mutation sites (CIMS), indicating a direct AGO:RNA association. In addition to the largest peak, which matches miR-29 (referred to here as the *Lep 1* binding site), a secondary miR-29 binding site (*Lep 2*) is located 731 bp distal to the first and was also validated by luciferase assay (Fig. 2i). Following normalization to leptin mRNA abundance, average tags per peak for both *Lep 1* and *Lep 2* were greater in iBAT, suggesting that miR-29 regulation contributes to suppressed leptin production in iBAT relative to eWAT (Supplementary Fig. 2d-f). The other peaks in the leptin 3’-UTR matched miRNA seed sites of miR-7, miR-125 and miR-30. The second tallest peak, corresponding to a potential miR-125 binding site, did not validate by luciferase assay (data not shown).

We next assessed whether miR-29 was capable of repressing leptin in primary adipocytes following transfection of a miR-29a mimic. 72 hours post-transfection, miR-29a levels were substantially increased in both iBAT and eWAT (Fig. 3a,d). Leptin mRNA levels correspondingly decreased by 40% in iBAT (p = 0.0054) and 80% in eWAT (p < 0.00001), relative to cells transfected with a mutated miR-29a mimic (Fig. 3b,e). Leptin secretion from primary adipocytes is lower than that from adipocytes *in vivo*; however, the conditioned media (CM) of primary cells contains a sufficient amount of leptin to be detected by ELISA. We observed a 50% decrease in leptin protein in CM from both iBAT and eWAT (Fig. 3c,f). As an alternative gain-of-function approach, primary cells were transduced with an adenovirus expressing miR-29a or a scrambled miRNA control. Consistent with the miRNA mimic experiments, an increase in miR-29a caused a 50–60% decrease in leptin mRNA in iBAT and eWAT (Supplementary Fig. 3a-d). Next, primary cells were transfected with a locked nucleic acid (LNA)^25^ to knock-down endogenous miR-29a. Both primary iBAT and eWAT adipocytes showed a significant decrease in miR-29a levels (Fig. 3g,j), and corresponding increases in leptin mRNA of 55% (p = 0.015) and 90% (p = 0.044), respectively (Fig. 3h,k). In CM from iBAT and eWAT, secreted leptin protein increased in parallel with leptin mRNA (Fig. 3i,l). For each experiment, RNA levels of the general adipocyte differentiation markers *Pparg2, Adipoq*, and *aP2* were measured. The expression of these differentiation dependent genes was moderately increased in iBAT and eWAT transfected with miRNA mimics (Supplementary Fig. 3e,f), consistent with previous reports that miR-29 promotes adipogenesis in human cells^26^. Adenoviral gain-of-function and LNA-based loss of function did not alter markers of adipocyte differentiation (Supplementary Fig. 3g-j).

**Fig. 3.**
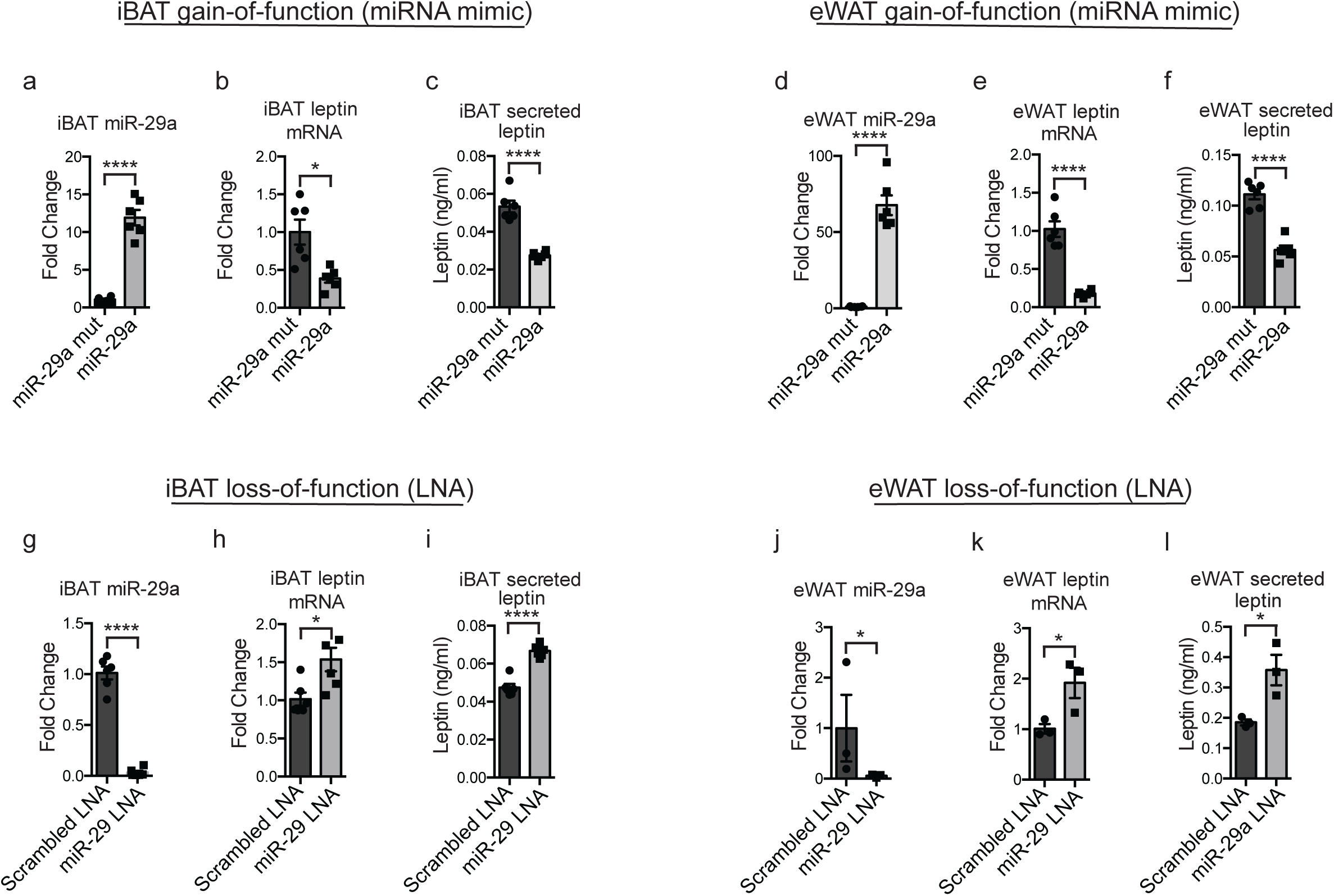
miR-29a Gain and loss-of-function modulates leptin levels *in vitro*. **a-c**, miR-29a (**a**), leptin mRNA (**b**), and CM leptin concentration (**c**) in primary iBAT cells transfected with a miR-29a mimic or a mutated miR-29 mimic (n = 6). **d-f**, miR-29a (**d**), leptin mRNA (**e**), and CM leptin concentration (**f**) in primary eWAT cells transfected with a miR-29a mimic or a mutated miR-29 mimic (n = 6). **g-i**, miR-29a (**g**), leptin mRNA (**h**), and CM leptin concentration (**i**) in primary iBAT cells transfected with a miR-29 LNA or a scrambled LNA (n = 6). **j-l**, miR-29a (**j**), leptin mRNA (**k**), and CM leptin concentration (**l**) in primary eWAT cells transfected with a miR-29 LNA or a scrambled LNA (n = 3). All results are presented as mean ±SEM. Student’s t test (*p < 0.05, ****p < 0.00005)

In obese mice and humans, leptin mRNA and circulating protein are generally increased compared to lean controls^27^, in proportion to body fat mass. We hypothesized that miR-29 levels would be inversely correlated with adiposity in obesity. Mice fed a HFD for 15 weeks are 60% heavier than their chow-fed counterparts (Fig. 4a) and have a 4-fold increase in leptin mRNA in iBAT, a 12-fold increase in eWAT, and an 8-fold increase in serum leptin (Fig. 4b,c). miR-29a levels were correspondingly reduced by approximately 30% in iBAT (p = 0.027) and 60% in eWAT (p < 0.00001) (Fig. 4d). To confirm our findings, we next measured miR-29 abundance in a genetic model of obesity. *Ob/ob* mice are hyperphagic and obese relative to littermate controls (Fig. 4e) due to a nonsense mutation in the leptin CDS that leads to high levels of leptin mRNA, but no functional protein (Fig. 4f,g). miR-29a in iBAT and eWAT of these mice was reduced by 24% (p = 0.017) and 31% (p = 0.011), respectively (Fig. 4h).

**Fig. 4:**
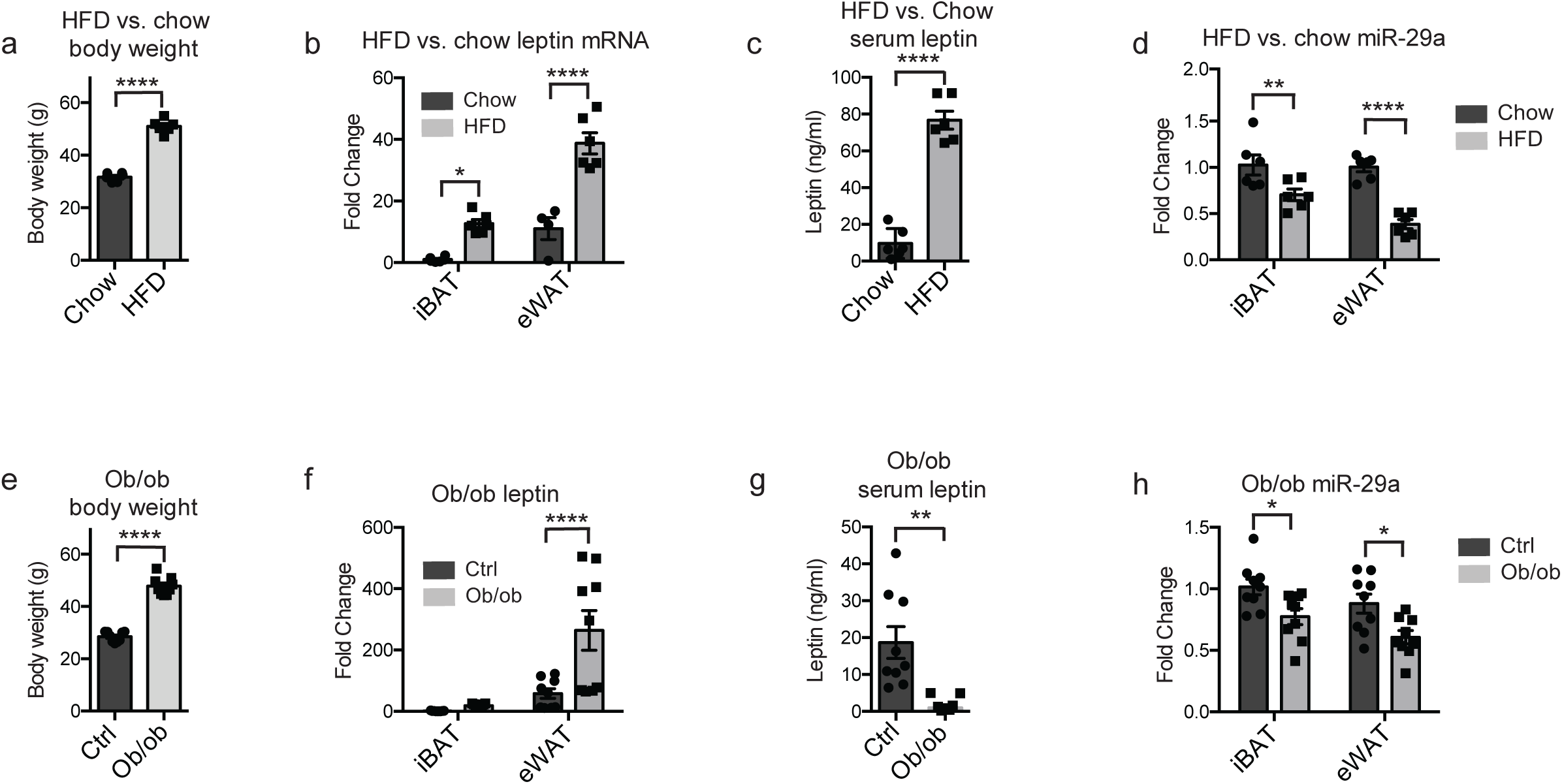
miR-29 abundance is inversely correlated with leptin in obese mice. **a-d**, Body weights (**a**), leptin mRNA (**b**), serum leptin (**c**) and miR-29a levels (**d**) in 21-week old HFD vs. chow mice (n = 6). **e-h**, Body weights (**e**), leptin mRNA (**f**), serum leptin (**g**) and miR-29a levels (**h**) in 9-week old ob^−/-^ vs. ob^+/-^ (n = 9). All results are presented as mean ±SEM. Student’s t test used for serum leptin comparisons and 2-way ANOVA used for leptin mRNA and miR-29a comparisons (*p < 0.05, **p < 0.005, ***p < 0.0005, ****p < 0.00005)

Human miR-29a is identical in sequence to murine miR-29a, and human leptin contains 2 TargetScan predicted binding sites^28^, analogous to those in mice. If miR-29a serves as a physiologically relevant regulator of leptin in humans, our findings suggest that miR-29a should be reduced in obese patients. Indeed, previously published work demonstrates that in a cohort of 66 men and women, ranging from lean to obese, miR-29a in visceral and subcutaneous WAT is negatively correlated with BMI^20^. A second human study found an increase in miR-29a in subcutaneous WAT from 19 obese individuals following diet and exercise-induced weight loss^29^.

In this study, we describe the first reported use of a high throughput, AGO crosslinking technique to map the miRNA targetome in adipose tissue. Our data indicate that abundant miRNAs associate with a vast array of targets, consistent with prior CLIP-based studies^12,30^. The miR-29 family was among the most active miRNAs, with over 1000 unique binding sites in iBAT and eWAT. Among the top-ranked miR-29 targets, we focused on leptin due to its central role in regulating energy homeostasis. Recent work has identified control of leptin transcription through an enhancer region 16 kb upstream of the transcriptional start site^31,32^, but until now, no post-transcriptional regulators of leptin have been described. This is surprising since miRNAs are estimated to regulate a majority of mRNAs in mammals^33^, and the leptin 3’-UTR is roughly 3-fold the length of an average 3’-UTR in vertebrates^34^. Our data show a substantial association between AGO and the leptin 3’-UTR, with 6 peaks reaching statistical significance, including 2 containing validated miR-29 binding sites.

Although miR-29 has not been studied extensively in adipose tissue, its role in a variety of cellular processes has been characterized previously. Our data suggest that many of these pathways may be similarly regulated in adipose tissue. In myocytes, miR-29 reduces insulin sensitivity by repressing *IRS-1*, a target which is also bound in adipose tissue^35^. HITS-CLIP also reveals miR-29 binding sites in *Myo1c* and *Cav2*, two genes involved in downstream propagation of insulin signaling^36,37^, suggesting overlapping function with the insulin de-sensitizing effect of miR-29 in liver^38,39^. In heart, miR-29 suppresses fibrosis by targeting *Col3a1, Fbn1*, and *Eln*^24^. These 3 genes, among several others including *Hspg2* and *Cflar*, are fibrosis-related miR-29 targets in adipose tissue^40,41^. Future *in vivo* studies, carried out in both lean and obese mice, are needed to fully explore the physiological role of miR-29 in these pathways.

Beyond miR-29, our HITS-CLIP data suggest a variety of other, previously unexplored roles for miRNAs in adipose tissue. The miR-101 family targets several genes involved in mitochondrial biogenesis, including *Gabpb2, Gabpa*, and *Atf2*. miR-101 also targets *Cidea*, a BAT enriched lipid-droplet associated protein^42^, and *Cebpa*, a transcription factor that controls adipogenesis^43^. miR-16 targets *Ogdh, Sucla2, Cox11, Srebp1, Ppara, Fasn*, and many other genes involved in the citric acid cycle, the electron transport chain, and cholesterol biosynthesis. miR-378, which has been previously characterized as a promoter of BAT expansion, binds the CDS of *UCP1* as its highest-ranked target. While seemingly paradoxical, this would be consistent with the reduction of *UCP1* in subcutaneous WAT seen in miR-378 transgenic mice^3^.

In conclusion, we have conducted the first comprehensive biochemical mapping of miRNA:mRNA binding in adipose tissue, providing an easily accessible resource for future studies. The HITS-CLIP results contain the breadth of data needed to describe miRNA activity at the network level, as well as the granularity to identify specific miRNA/target interactions. We have characterized miR-29 as one of the most active miRNAs in adipose tissue and validated leptin as one of miR-29’s top targets. As the only identified post-transcriptional regulator of leptin, miR-29 presents a unique target for therapeutic interventions aimed at treating obesity.

---

**Supplementary Fig. 1.**
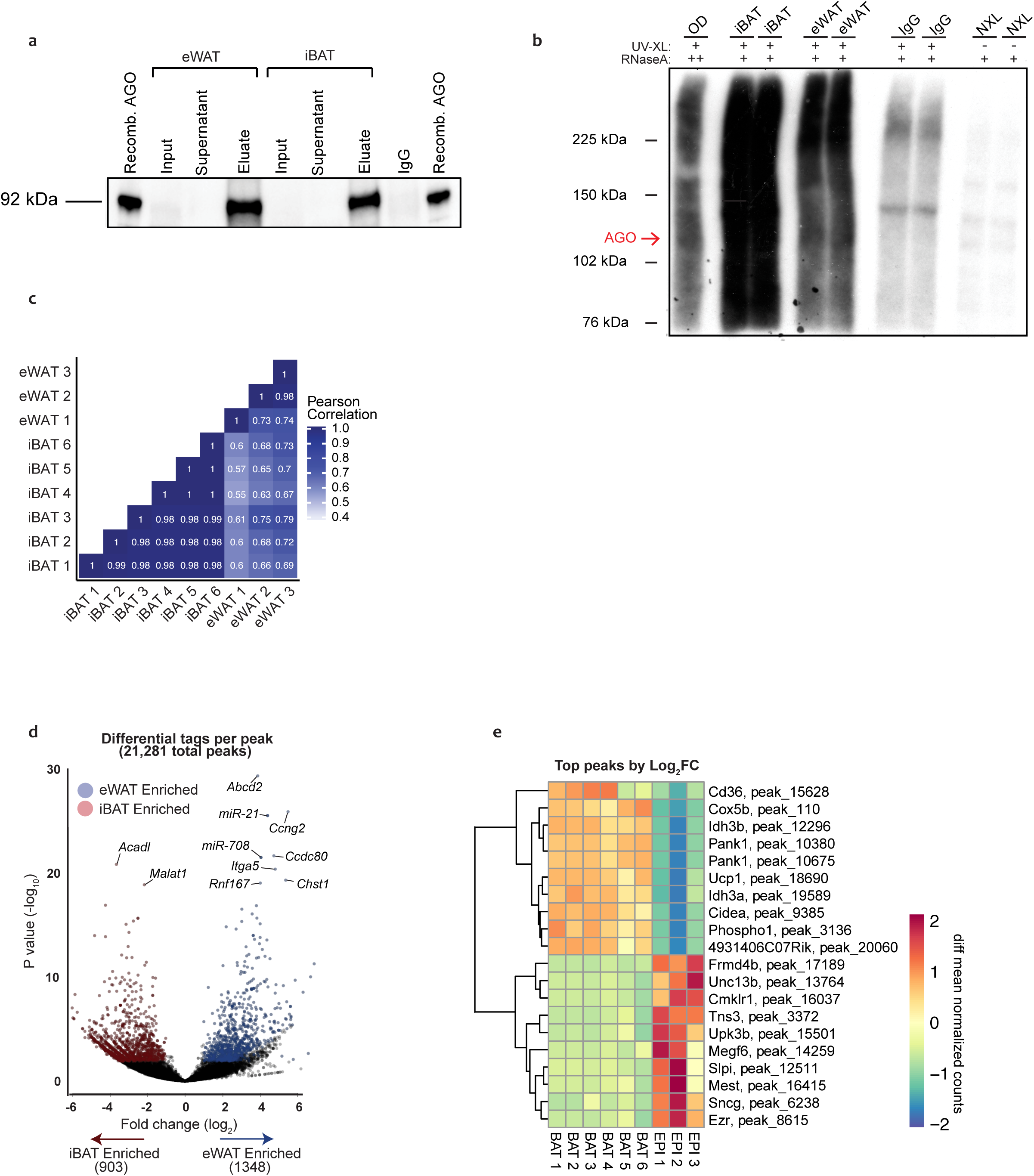
miRNA binding sites are variable across iBAT and eWAT. **a**, Immunoprecipitation of Ago from eWAT and iBAT tissues (image cropped to display AGO band more clearly). **b**, Autoradiogram showing NuPAGE separation of radio-labeled Ago-mRNA complexed, over-digested control (OD), iBAT, eWAT, an IgG control and a no-crosslink control (NXL). **c**, Pearson correlations for 6 iBAT and 3 eWAT samples. **d**, Volcano plot of iBAT vs. eWAT differential analysis of tags per peak. **e**, Heatmap of top 10 iBAT enriched and top 10 eWAT enriched peaks by tag abundance. Differential analysis of tags performed using DESEQ2 with a significance cutoff of FDR < 0.05 and |Log_2_FC| > 1.

**Supplementary Fig. 2.**
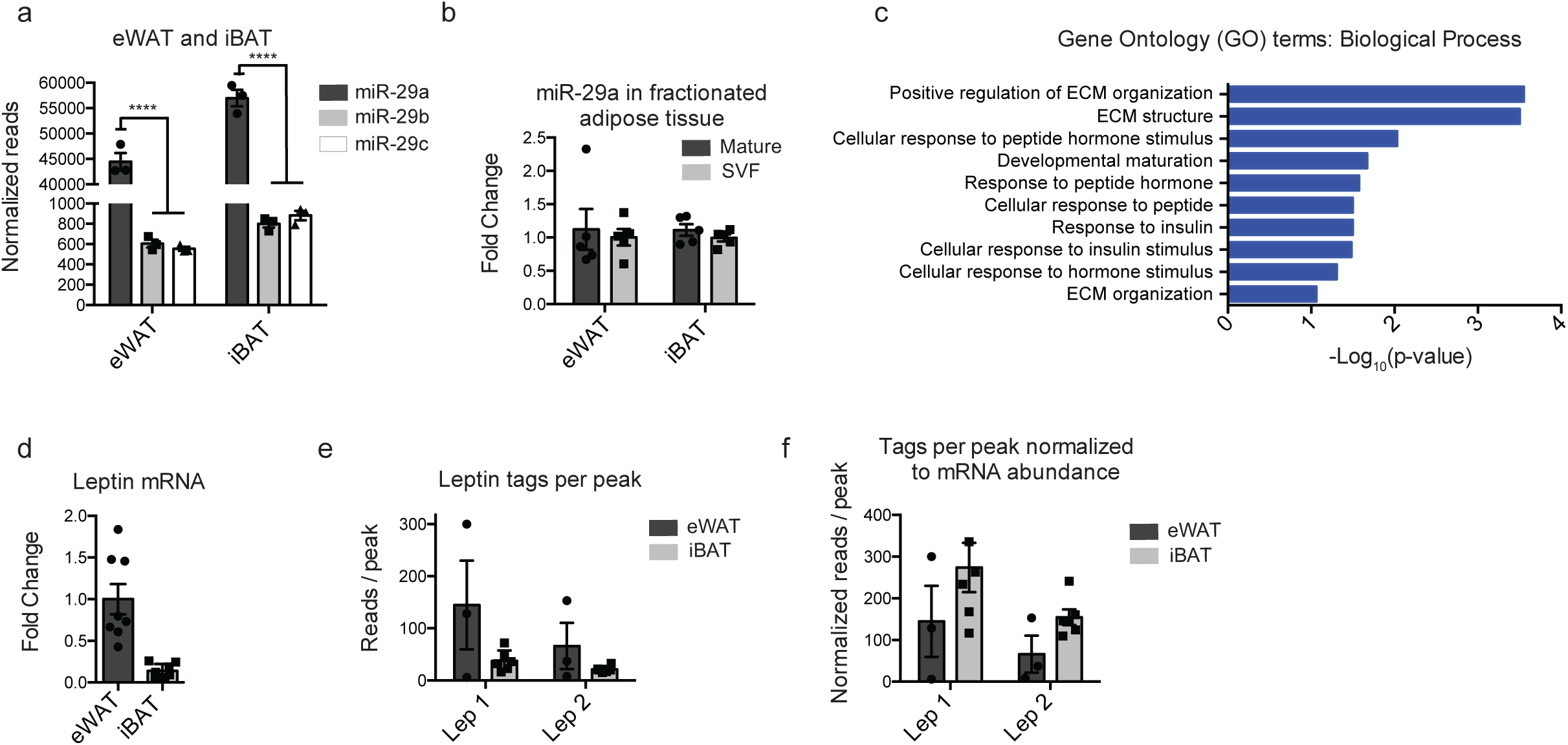
Leptin transcripts are more likely to be bound by miR-29 in iBAT than eWAT. **a**, DEseq normalized tag abundance of miR-29 from iBAT and eWAT of 13-week old male mice (n = 3). **b**, miR-29a levels in digested and fractionated iBAT and eWAT (n = 5). **c**, Gene ontology term enrichment of top 250 miR-29 3’-UTR targets. **d**, Leptin mRNA abundance in iBAT vs eWAT from 15-week old male mice (n = 8). **e,f** Average tags per peak in *Lep 1* and *Lep 2* binding sites, unnormalized (**e**) and normalized to leptin mRNA abundance (**f**) (2 mice pooled per group, n = 6 for BAT, n = 3 for eWAT). All results are presented as mean ±SEM. Student’s t test for eWAT vs. iBAT in (A) (***p < 0.0005, ****p < 0.00005)

**Supplementary Fig. 3.**
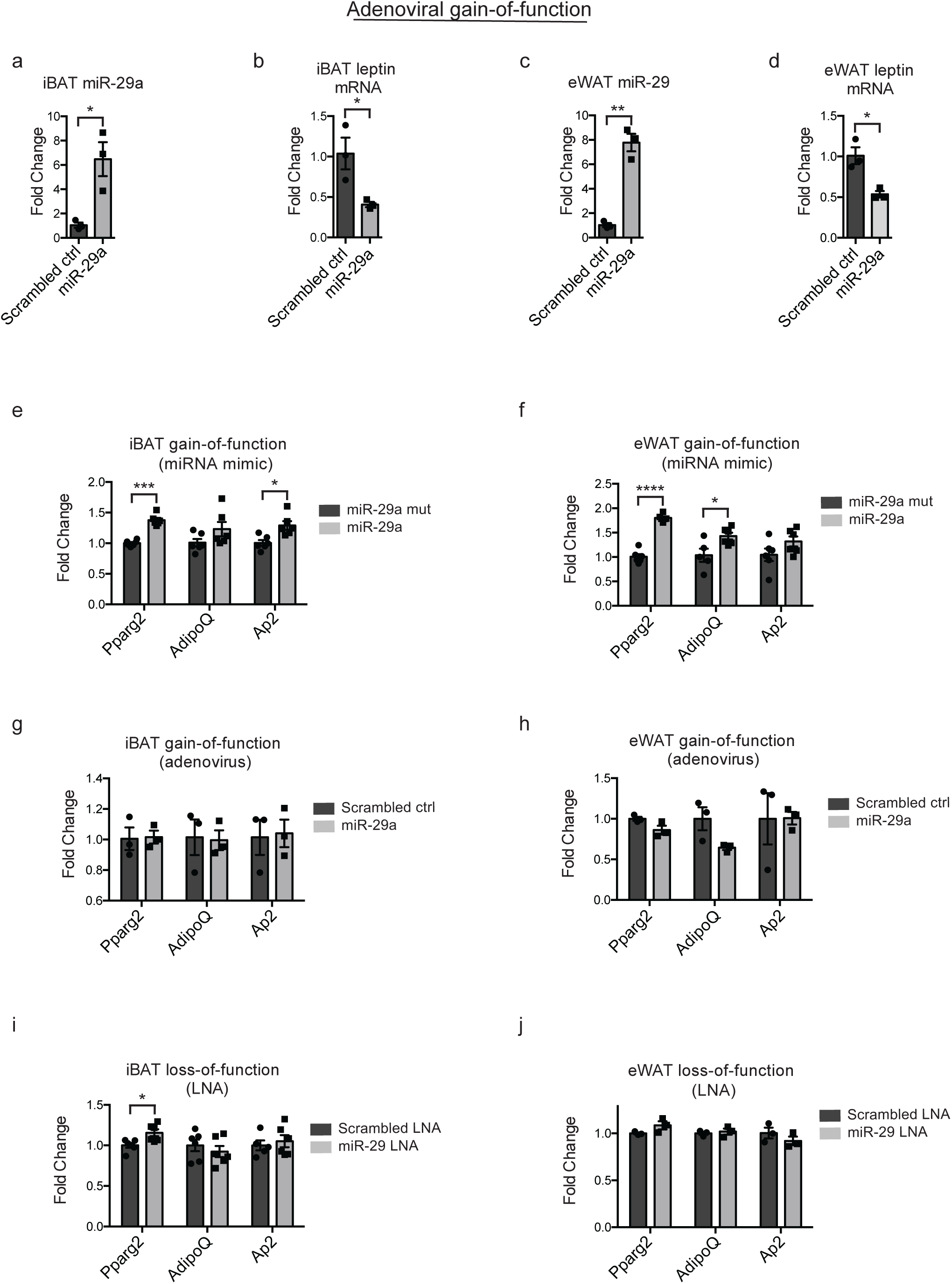
miR-29a adenoviral gain-of-function represses leptin mRNA *in vitro*. **a,b**, miR-29a (**a**) and leptin mRNA (**b**) in primary iBAT cells transduced with adenovirus containing primary miR-29a or a scrambled sequence (n = 3). **c,d**, miR-29a (**c**) and leptin mRNA (**d**) in primary eWAT cells transduced with adenovirus containing primary miR-29a or a scrambled sequence (n = 3). **e,j**, qPCR of differentiation markers in primary iBAT and eWAT transfected with miR-29a mimic (**e,f**), transduced with miR-29a adenovirus (**g,h**) or transfected with miR-29 LNA (**i,j**) (n = 3 for **e, f** and **g**, n = 6 for **h, i**, and **j**). All results are presented as mean ±SEM. Student’s t test for gain/loss of function vs. control (*p < 0.05, **p < 0.005, ****p < 0.00005)

## METHODS

### Mouse models

Animal care and experimentation were performed according to procedures approved by the Institutional Animal Care and Use Committee at the Rockefeller University. All mice used in the study were purchased from The Jackson Laboratory and were inbred on the C57BL/6 background, including *ob/ob* (stock no. 000632), Wt (stock no. 000664) and HFD (stock no. 380050). All mice used were male. Controls for *ob/ob* mice were heterozygous littermates. Mice were group housed at 5 mice per cage at 23° C and maintained on 12-hour light:dark cycles. HFD mice were fed a diet composed of 60% dietary fat (Research Diets Cat# D12492) from 6 weeks of age, and all other animals were maintained on a chow diet, ad libitum.

### HEK-293A cell line

HEK-293A cells (Invitrogen, Cat#R70507) were maintained at 37°C with 5% CO_2_ in DMEM (4.5 g/L glucose, 0.58 g/L L-glutamine, 110 mg/L sodium pyruvate) supplemented with 10% FBS (Gemini Bio-Products) and 1% penicillin-streptomycin (Gibco).

### Primary adipocyte culture

Primary adipocytes were isolated from 6-week old male C57BL/6J eWAT and iBAT adipose depots. Dissected adipose depots were incubated in 10 mL digestion buffer in a 37°C shaking water bath (140 RPM) for 16 minutes (eWAT) or 40 minutes (iBAT). iBAT digestion buffer was prepared as follows: 15 mg collagenase B (Roche, Cat#11088831001) was dissolved in 5 mL of 2X master mix consisting of 125 mM NaCl, 5 mM KCl, 1.3 mM CaCl_2_, 5 mM glucose, 1% penicillin-streptomycin, 4% BSA in H_2_O and 5 mL PBS. eWAT digestion buffer was prepared as follows: 100 mg collagenase D (Roche, Cat#11088882001) was dissolved in a solution of 10 mL PBS, 200 μL of 120 mg/mL dispase II (Roche, Cat#04942078001) and 40 μL 2.5 M CaCl_2_. Following digestion, the stromal vascular fraction was isolated by centrifugation and filtration at 100 μm and 70 μm, then plated in 12-well collagen coated plates.

All primary adipocyte cultures were maintained at 37°C with 10% CO_2_. Prior to differentiation, iBAT cells were cultured in GlutaMAX media (Gibco, Cat#10565) supplemented with 10% FBS and 1% penicillin-streptomycin) and eWAT cells were cultured in ITS media (ITS media: 56.65% 1g/L glucose DMEM (Gibco, Cat#11885-084), 37.65% 1X MCDB210 pH 7.25 (Sigma, Cat#M6770), 2% FBS, 2% ITS premix (Corning, Cat#354352), 1% 10 mM L-ascorbic acid 2-phosphate in DMEM, 0.01% 100 μg/mL bFGF (BD Bio, Cat#13256-029), 0.5% penicillin-streptomycin, and 0.2% primocin). Following induction of differentiation, iBAT cells and eWAT cells were cultured in GlutaMAX media.

When iBAT preadipocytes reached 100% confluency and eWAT preadipocytes reached 95% confluency, differentiation was induced using a standard cocktail of 0.5 mM IBMX, 1 mM dexamethasone, 850 nM insulin, and 1 mM rosiglitazone dissolved in GlutaMAX media. After 48 hours, media was replaced with fresh media containing only insulin and rosiglitazone, and after 96 hours media was replaced with fresh media containing only insulin.

### Adenovirus cloning, amplification and transduction

Adenoviruses for miR-29a and the scrambled miRNA control were cloned using the AdEasy system (Agilent, Cat#240010) according to the protocol provided. Briefly, primary miRNAs for miR-29a and miR-16 were amplified from genomic DNA by PCR and cloned into the pAdTrack shuttle (Addgene, Plasmid#16404). The miR-16 pAdTrack plasmid was then mutated by site-directed mutagenesis to induce 3 point mutations into the miR-16 seed sequence, creating the scrambled control. Both the miR-29a and the scrambled control pAdTrack plasmids were linearized with PmeI and transformed into BJ5183-AD-1 competent cells for recombination with the adenoviral backbone. Successful clones were amplified in the XL10-Gold bacterial strain. Generation of the initial adenoviral stock was performed by digesting the adenoviral plasmid with PacI, then transfecting HEK-293A cells. Several rounds of amplification were performed by infecting HEK-293A cells with low titer adenovirus. For transduction of primary adipocytes, iBAT and eWAT cells were treated with a final adenoviral concentration of 7 x 10^7^ IFU/mL on the 4^th^ day of differentiation. 72 hours post-transduction, adipocytes were inspected for GFP expression, then total RNA was collected.

### HITS-CLIP Protocol

iBAT and eWAT tissues were dissected from 15-week old C57BL/6J mice and immediately flash frozen in liquid nitrogen. Frozen iBAT and eWAT pairs were then pooled and ground while still frozen with a Cellcrusher tissue pulverizer (Cellcrusher) cooled in liquid nitrogen. Frozen samples were spread to a thin layer on petri dishes pre-cooled on dry ice and UV-crosslinked 3 times at 400 mJ/cm^2^ using a Stratalinker 2400 (Stratagene). All samples were stored at −80°C until tissue lysis.

For each sample, 400 μL Protein A Dynabeads (Invitrogen) were pooled, then prepared by washing with 1 mL antibody binding (AB) buffer (AB: PBS, 0.02% tween 20) 3 times. Following the final wash, beads were resuspended in 400 μL AB per sample and 50 μL of rabbit anti-mouse bridging antibody (Jackson ImmunoResearch, Cat#315005008) was added. Beads were rotated at 22°C for 1 hour, washed 3 times with 1 mL AB, resuspended in AB, and incubated for 2 hours at 22°C with either 12 μL pan-Ago antibody (Millipore clone 2A8^47^, manufactured at Rockefeller), or 12 μL mouse normal IgG (Santa Cruz Biotechnology, Cat#sc-2343).

Tissue pellets were lysed with 800μl PXL+ lysis buffer (PXL+: 1x PBS; 0.1% SDS; 0.5% Na-DOC; 0.5% NP-40, one Complete EDTA-Free Protease Inhibitor Cocktail tablet (Roche) per 10 mL) and subject to DNase (Promega, Cat#M610A) treatment by incubating for 5 minutes at 37°C, shaking at 1100 rpm. Samples were then centrifuged at 20,000 x g for 20 minutes, and a needle was used to withdraw 800 μL of aqueous protein lysate while avoiding the lipid layer and pellet. Lysates were then treated with 20U/mL RNase A (Affymetrix, discontinued) by first diluting the RNAse to the indicated concentration by volume (1:100 for “High RNase,” or 1:10,000 for “Low RNase”) then treating samples with 8μl diluted RNAse per mL of lysate and digesting for 5 minutes at 37°C, 1100 rpm. Immediately following RNAse digestion, samples were added to prepared antibody bound beads and rotated for 3 hours at 4°C.

After IP, the following washes were performed: twice with lysis buffer, twice with high salt lysis buffer (5X PBS, 1% Igepal, 0.5% deoxycholate and 0.1% SDS), twice with stringent wash buffer (15 mM Tris pH 7.5, 5 mM EDTA, 2.5 mM EGTA, 1% TritonX-100, 1% NaDOC, 0.1% SDS, 120 mM NaCl, 25 mM KCl), twice with high salt wash buffer (15 mM Tris pH 7.5, 5 mM EDTA, 2.5 mM EGTA, 1% TritonX-100, 1% NaDOC, 0.1% SDS, 1M NaCl), twice with low salt wash buffer (15 mM Tris pH 7.5, 5 mM EDTA), and twice with PNK wash buffer (50 mM Tris pH 7.4, 10 mM MgCl2, 0.5% NP-40). For the second of each wash, specimens were rotated for 2-3 minutes at 22°C. Tags were dephosphorylated as described^47^ and subject to overnight 3’ ligation at 16°C with a pre-adenylated linker^48^ with the following ligation reaction: 2 μL of 25 μM linker, 2 μL of T4 RNA Ligase 2, truncated K227Q (NEB), 1X ligation buffer (supplied with ligase), 2 μL Superasin RNase inhibitor (Promega), and 8 μL PEG8000 (supplied with ligase). The resulting beads were washed, ^32^P-labeled, and subject to SDS-PAGE and transfer as described^47^.

Tags were collected from nitrocellulose as described^49^ with the following exceptions: Phenol:Chloroform:IAA, 25:24:1 pH 6.6 was used for extraction, and tags were precipitated with a standard NaOAC precipitation. Cloning was performed using the BrdU-CLIP protocol as described^48^ with a few exceptions. Briefly, the RT primer contains a 14-nt degenerate linker (a 3-nt degenerate sequence, a 4-nt multiplexing index, and a 7-nt unique molecular identifier), a 5′linker for PCR amplification, a spacer to prevent rolling circle amplification after circularization, and the reverse-complementary sequence of the 3′linker for reverse transcription. BrdUTP-labeled cDNA was specifically isolated via two sequential BrdUTP immunoprecipitations (Abcam AB8955) and circularized with CircLigase II (Epicenter CL9025K). Ago-CLIP libraries were sequenced by MiSeq (Illumina) to obtain 75-nt single end reads.

### Western blot analysis

Samples and beads were prepared as described above. Following immunoprecipitation and washes, samples were immediately eluted, separated by SDS-PAGE, transferred to a nitrocellulose membrane and incubated with anti-Ago 2A8. The Ago positive control was prepared from cell lysate of HEK-293T cells transfected with a plasmid expressing mouse Ago2. Protein was detected by incubation with HRP-conjugated sheep anti-mouse secondary antibody (Jackson ImmunoResearch) and blots were imaged on a Biorad Chemidoc Imaging System.

### CLIP Tool Kit (CTK) analysis of HITS-CLIP data

Sequencing files were demultiplexed (barcode.txt), linkers were trimmed and reads were collapsed. Reads were aligned to mm10 genome assembly using Novoalign. All raw CLIP read processing and significant peak calling was performed using the CTK software package^50^. CLIP peaks and CIMS analysis was carried out as previously described^47^.

### Analysis performed in R

CLIP peak and unique tag output files were analyzed in R^51^ using Bioconductor^52^. using packages, “AnnotationDbi”, “annotatr”, “Biobase”, “BiocGenerics”, “BiocManager”, “BiocParallel”, “Biostrings”, “BSgenome”, “BSgenome.Mmusculus.UCSC.mm10”, “DelayedArray”, “DESeq2”, “dplyr”, “edgeR”, “GenomeInfoDb”, “GenomicFeatures”, “GenomicRanges”, “ggplot2”, “ggpubr”, “ggsignif”, “gridExtra”, “IRanges”, “limma”, “magrittr”, “matrixStats”, “mirbase.db”, “org.Mm.eg.db”, “plyr”, “Rsamtools”, “rtracklayer”, “S4Vectors”, “stringr”, “SummarizedExperiment”, “tibble”, “tidyr”, “TxDb.Mmusculus.UCSC.mm10.knownGene”, “tximport”, “VariantAnnotation”, “wesanderson”, “XVector”. R version 3.6.1 Patched (2019-08-29 r77095).

### AGO-associated miRNA mapping and analysis

All HITS-CLIP reads were mapped to a catalog of mature murine miRNAs from miRbase, and miRNAs that were not detected in adipose tissue by small RNAseq were discarded. Next, reads were normalized by RPM, then grouped into annotated families. miRNAs were then matched to peaks by scanning for 6-mer, 7-mer (including 1-7 and 2-8 matching base pairs) and 8-mer seed sequences among peaks, limited to miRNAs with average normalized reads > 10.

### Dual-luciferase assays

5’-phosphorylated oligonucleotides containing the predicted binding sites of miR-29 target genes, 6 bp of flanking DNA, and SacI/XhoI restriction site overhands were annealed then ligated into the dual luciferase reporter plasmid (Promega, Cat#E1960). For mutated sequences, 3-4-points were induced in the miR-29a seed binding site using the Q5 Site Directed Mutagenesis Kit (NEB, Cat#E0554S). 20 ng of recombinant dual-luciferase plasmid and 6 pmol of either miR-29a mimic or scrambled control were mixed with 100 μL Opti-MEM containing 1 μL lipofectamine RNAiMax (Invitrogen, Cat.#13778) in 24-well plate wells. Reverse transfection was initiated by adding 90,000 HEK-293A cells/well in 500 μL. After 48 hours, firefly activity was measured by luminescence and normalized to Renilla activity.

### LNA and miRNA mimic transfections

All cells were transfected on 12-well plates 4 days after inducing differentiation. For each well, 3 μL of 10 μM double-stranded mature miRNA mimic (Dharmacon) or 20 μM LNA (Qiagen) was mixed with 3 μL lipofectamine RNAiMAX in 100 μL Opti-MEM reduced serum media and added to cells following a 15-minute incubation. RNA was collected in TRIzol (Invitrogen) 72 hours after transfection.

### RNA extraction, reverse transcriptase PCR, and quantitative PCR

RNA was extracted from tissues and cells using a TRIzol (Invitrogen)/chloroform extraction, followed by purification with RNeasy mini kits (Qiagen). To retain small RNAs during RNA purification, 1.5X volumes of 100% ethanol was used to precipitate RNA prior to column loading and RW1 buffer was substituted with RWT (Qiagen). For cDNA synthesis of mRNAs, the Applied Biosystems high capacity cDNA synthesis kit was used with 1 μg of purified RNA. For cDNA synthesis of miRNAs, reverse transcription using the universal primer 5’ - CAGGTCCAGTTTTTTTTTTTTTTTVN - 3’ was used as previously reported^53^. qPCR for both mRNAs and miRNAs was performed on a QuantStudio 6 Flex machine (Applied Biosystems) using SYBR-green fluorescent dye (Applied Biosystems). qPCR reactions were carried out in 384-well plates with a volume of 10 μL containing 0.5 μM forward and reverse primers, 1X qPCR master mix and 5 ng of mRNA-based cDNA or 1 ng of miRNA-based cDNA. MiRprimer2.0 was used to design primers for miRNA qPCR^54^.

### Leptin ELISA

A mouse leptin ELISA kit (Crystal Chem, Cat#90030) was used to measure leptin concentrations in serum (5 μL per assay) or CM (100 μL per assay) according to manufacturer’s instructions.

### Adipose tissue fractionation

Dissected adipose depots were incubated in 10 mL digestion buffer in a 37°C shaking water bath (140 RPM) for 16 minutes (eWAT) or 40 minutes (iBAT). Following digestion, the stromal vascular fraction and mature adipocyte fraction were separated by centrifugation (10 minutes at 500 x g). Fractions were added directly to TRIzol (Invitrogen, Cat#15596026) for total RNA extraction.

### Notes on statistical analyses

All measurements and analyses were performed using distinct samples. All t-tests were two-tailed with an assumption of normality. Significance cut-offs for HITS-CLIP tag differential comparisons (tags per gene and tags per peak) were adjusted for multiple comparisons.

## DATA AVAILABILITY

HITS-CLIP raw data are available through Gene Expression Omnibus (GSE142677). All other data, including raw data, are available upon reasonable request from the corresponding author.

## CODE AVAILABILITY

The code used for the HITS-CLIP analysis and the workflow used on Galaxy are available upon reasonable request from the corresponding author.

## ACKNOWLEDGEMENTS

This work was supported by The Rockefeller University Sackler Center for Biomedicine and Nutrition Research and by the American Diabetes Association Pathway to Stop Diabetes Grant 1-17-ACE-17 (PC). S.K.S. was supported by a Medical Scientist Training Program grant from the National Institute of General Medical Sciences of the National Institutes of Health under award number T32GM007739 to the Weill Cornell/Rockefeller/Sloan Kettering Tri-Institutional MD-PhD Program. We would like to thank C.H.J. Choi for providing samples from lean and obese mice and J. Chi for technical assistance with primary adipocyte cultures. We would also like to thank T.S. Carroll for providing the code used to calculate PhastCons scores.

## AUTHOR CONTRIBUTIONS

S.O., P.C. and E.M. designed and planned the study and R.B.D. and P.S. provided technical and strategic input. E.M. and S.O. carried out the HITS-CLIP experiments and analysis, S.O. and S.S. performed luciferase assays, primary cell gain-and-loss of function experiments, and mouse studies. S.O., E.M., M.K., and F.M. carried out bioinformatic analyses. The manuscript was written by S.O. and P.C. with contributions from all authors.

## REFERENCES

1 Mori, M., Nakagami, H., Rodriguez-Araujo, G., Nimura, K. & Kaneda, Y. Essential role for miR-196a in brown adipogenesis of white fat progenitor cells. PLoS Biol 10, e1001314, doi:10.1371/journal.pbio.1001314 (2012).

2 Trajkovski, M., Ahmed, K., Esau, C. C. & Stoffel, M. MyomiR-133 regulates brown fat differentiation through Prdm16. Nat Cell Biol 14, 1330–1335, doi:10.1038/ncb2612 (2012).

3 Pan, D. et al. MicroRNA-378 controls classical brown fat expansion to counteract obesity. Nat Commun 5, 4725, doi:10.1038/ncomms5725 (2014).

4 Cowley, M. A. et al. Leptin activates anorexigenic POMC neurons through a neural network in the arcuate nucleus. Nature 411, 480–484, doi:10.1038/35078085 (2001).

5 Tontonoz, P. & Spiegelman, B. M. Fat and beyond: the diverse biology of PPARgamma. Annu Rev Biochem 77, 289–312, doi:10.1146/annurev.biochem.77.061307.091829 (2008).

6 Seale, P. et al. Transcriptional control of brown fat determination by PRDM16. Cell Metab 6, 38–54, doi:10.1016/j.cmet.2007.06.001 (2007).

7 Cohen, P. et al. Ablation of PRDM16 and beige adipose causes metabolic dysfunction and a subcutaneous to visceral fat switch. Cell 156, 304–316, doi:10.1016/j.cell.2013.12.021 (2014).

8 Mori, M. A. et al. Altered miRNA processing disrupts brown/white adipocyte determination and associates with lipodystrophy. J Clin Invest 124, 3339–3351, doi:10.1172/JCI73468 (2014).

9 Reis, F. C. et al. Fat-specific Dicer deficiency accelerates aging and mitigates several effects of dietary restriction in mice. Aging (Albany NY) 8, 1201–1222, doi:10.18632/aging.100970 (2016).

10 Acharya, A. et al. miR-26 suppresses adipocyte progenitor differentiation and fat production by targeting Fbxl19. Genes Dev 33, 1367–1380, doi:10.1101/gad.328955.119 (2019).

11 Luna, J. M. et al. Argonaute CLIP Defines a Deregulated miR-122-Bound Transcriptome that Correlates with Patient Survival in Human Liver Cancer. Mol Cell 67, 400–410 e407, doi:10.1016/j.molcel.2017.06.025 (2017).

12 Chi, S. W., Zang, J. B., Mele, A. & Darnell, R. B. Argonaute HITS-CLIP decodes microRNA-mRNA interaction maps. Nature 460, 479–486, doi:10.1038/nature08170 (2009).

13 Spengler, R. M. et al. Elucidation of transcriptome-wide microRNA binding sites in human cardiac tissues by Ago2 HITS-CLIP. Nucleic Acids Res 44, 7120–7131, doi:10.1093/nar/gkw640 (2016).

14 Siepel, A. et al. Evolutionarily conserved elements in vertebrate, insect, worm, and yeast genomes. Genome Res 15, 1034–1050, doi:10.1101/gr.3715005 (2005).

15 Gabriely, I. et al. Removal of visceral fat prevents insulin resistance and glucose intolerance of aging: an adipokine-mediated process? Diabetes 51, 2951–2958, doi:10.2337/diabetes.51.10.2951 (2002).

16 Ventayol, M. et al. miRNA let-7e targeting MMP9 is involved in adipose-derived stem cell differentiation toward epithelia. Cell Death Dis 5, e1048, doi:10.1038/cddis.2014.2 (2014).

17 Giroud, M. et al. Let-7i-5p represses brite adipocyte function in mice and humans. Sci Rep 6, 28613, doi:10.1038/srep28613 (2016).

18 Hu, F. et al. miR-30 promotes thermogenesis and the development of beige fat by targeting RIP140. Diabetes 64, 2056–2068, doi:10.2337/db14-1117 (2015).

19 Koh, E. H. et al. miR-30a Remodels Subcutaneous Adipose Tissue Inflammation to Improve Insulin Sensitivity in Obesity. Diabetes 67, 2541–2553, doi:10.2337/db17-1378 (2018).

20 Glantschnig, C. et al. A miR-29a-driven negative feedback loop regulates peripheral glucocorticoid receptor signaling. FASEB J 33, 5924–5941, doi:10.1096/fj.201801385RR (2019).

21 Roderburg, C. et al. Micro-RNA profiling reveals a role for miR-29 in human and murine liver fibrosis. Hepatology 53, 209–218, doi:10.1002/hep.23922 (2011).

22 Toyono, T. et al. MicroRNA-29b Overexpression Decreases Extracellular Matrix mRNA and Protein Production in Human Corneal Endothelial Cells. Cornea 35, 1466–1470, doi:10.1097/ICO.0000000000000954 (2016).

23 Li, J. et al. miR-29b contributes to multiple types of muscle atrophy. Nat Commun 8, 15201, doi:10.1038/ncomms15201 (2017).

24 van Rooij, E. et al. Dysregulation of microRNAs after myocardial infarction reveals a role of miR-29 in cardiac fibrosis. Proc Natl Acad Sci U S A 105, 13027–13032, doi:10.1073/pnas.0805038105 (2008).

25 Grunweller, A. & Hartmann, R. K. Locked nucleic acid oligonucleotides: the next generation of antisense agents? BioDrugs 21, 235–243, doi:10.2165/00063030-200721040-00004 (2007).

26 Zhang, X. M. et al. MicroRNA-29b promotes the adipogenic differentiation of human adipose tissue-derived stromal cells. Obesity (Silver Spring) 24, 1097–1105, doi:10.1002/oby.21467 (2016).

27 Frederich, R. C. et al. Leptin levels reflect body lipid content in mice: evidence for diet-induced resistance to leptin action. Nat Med 1, 1311–1314, doi:10.1038/nm1295-1311 (1995).

28 Agarwal, V., Bell, G. W., Nam, J. W. & Bartel, D. P. Predicting effective microRNA target sites in mammalian mRNAs. Elife 4, doi:10.7554/eLife.05005 (2015).

29 Kristensen, M. M. et al. miRNAs in human subcutaneous adipose tissue: Effects of weight loss induced by hypocaloric diet and exercise. Obesity (Silver Spring) 25, 572–580, doi:10.1002/oby.21765 (2017).

30 Hafner, M. et al. Transcriptome-wide identification of RNA-binding protein and microRNA target sites by PAR-CLIP. Cell 141, 129–141, doi:10.1016/j.cell.2010.03.009 (2010).

31 Dallner, O. S. et al. Dysregulation of a long noncoding RNA reduces leptin leading to a leptin-responsive form of obesity. Nat Med 25, 507–516, doi:10.1038/s41591-019-0370-1 (2019).

32 Zhang, Y. et al. A noncanonical PPARgamma/RXRalpha-binding sequence regulates leptin expression in response to changes in adipose tissue mass. Proc Natl Acad Sci U S A 115, E6039–E6047, doi:10.1073/pnas.1806366115 (2018).

33 Friedman, R. C., Farh, K. K., Burge, C. B. & Bartel, D. P. Most mammalian mRNAs are conserved targets of microRNAs. Genome Res 19, 92–105, doi:10.1101/gr.082701.108 (2009).

34 Mignone, F. & Pesole, G. mRNA Untranslated Regions (UTRs). eLS, 1–6, doi:10.1002/9780470015902.a0005009.pub3 (2018).

35 Yang, W. M., Jeong, H. J., Park, S. Y. & Lee, W. Induction of miR-29a by saturated fatty acids impairs insulin signaling and glucose uptake through translational repression of IRS-1 in myocytes. FEBS Lett 588, 2170–2176, doi:10.1016/j.febslet.2014.05.011 (2014).

36 Bose, A. et al. Glucose transporter recycling in response to insulin is facilitated by myosin Myo1c. Nature 420, 821–824, doi:10.1038/nature01246 (2002).

37 Kwon, H., Lee, J., Jeong, K., Jang, D. & Pak, Y. Fatty acylated caveolin-2 is a substrate of insulin receptor tyrosine kinase for insulin receptor substrate-1-directed signaling activation. Biochim Biophys Acta 1853, 1022–1034, doi:10.1016/j.bbamcr.2015.02.002 (2015).

38 Hung, Y. H. et al. Acute suppression of insulin resistance-associated hepatic miR-29 in vivo improves glycemic control in adult mice. Physiol Genomics 51, 379–389, doi:10.1152/physiolgenomics.00037.2019 (2019).

39 Kurtz, C. L. et al. MicroRNA-29 fine-tunes the expression of key FOXA2-activated lipid metabolism genes and is dysregulated in animal models of insulin resistance and diabetes. Diabetes 63, 3141–3148, doi:10.2337/db13-1015 (2014).

40 Lord, M. S. et al. The multifaceted roles of perlecan in fibrosis. Matrix Biol 68–69, 150-166, doi:10.1016/j.matbio.2018.02.013 (2018).

41 McCubbrey, A. L. et al. Deletion of c-FLIP from CD11b(hi) Macrophages Prevents Development of Bleomycin-induced Lung Fibrosis. Am J Respir Cell Mol Biol 58, 66–78, doi:10.1165/rcmb.2017-0154OC (2018).

42 Barneda, D. et al. The brown adipocyte protein CIDEA promotes lipid droplet fusion via a phosphatidic acid-binding amphipathic helix. Elife 4, e07485, doi:10.7554/eLife.07485 (2015).

43 Wang, N. D. et al. Impaired energy homeostasis in C/EBP alpha knockout mice. Science 269, 1108–1112, doi:10.1126/science.7652557 (1995).

44 Zhou, S. et al. MicroRNA-29b-3p Targets SPARC Gene to Protect Cardiocytes against Autophagy and Apoptosis in Hypoxic-Induced H9c2 Cells. J Cardiovasc Transl Res 12, 358–365, doi:10.1007/s12265-018-9858-1 (2019).

45 Chen, T. et al. MicroRNA-29a regulates pro-inflammatory cytokine secretion and scavenger receptor expression by targeting LPL in oxLDL-stimulated dendritic cells. FEBS Lett 585, 657–663, doi:10.1016/j.febslet.2011.01.027 (2011).

46 Qiang, J., Tao, Y. F., He, J., Sun, Y. L. & Xu, P. miR-29a modulates SCD expression and is regulated in response to a saturated fatty acid diet in juvenile genetically improved farmed tilapia (Oreochromis niloticus). J Exp Biol 220, 1481–1489, doi:10.1242/jeb.151506 (2017).

47 Moore, M. J. et al. Mapping Argonaute and conventional RNA-binding protein interactions with RNA at single-nucleotide resolution using HITS-CLIP and CIMS analysis. Nat Protoc 9, 263–293, doi:10.1038/nprot.2014.012 (2014).

48 Moore, M. J. et al. ZFP36 RNA-binding proteins restrain T cell activation and anti-viral immunity. Elife 7, doi:10.7554/eLife.33057 (2018).

49 Zarnegar, B. J. et al. irCLIP platform for efficient characterization of protein-RNA interactions. Nat Methods 13, 489–492, doi:10.1038/nmeth.3840 (2016).

50 Shah, A., Qian, Y., Weyn-Vanhentenryck, S. M. & Zhang, C. CLIP Tool Kit (CTK): a flexible and robust pipeline to analyze CLIP sequencing data. Bioinformatics 33, 566–567, doi:10.1093/bioinformatics/btw653 (2017).

51 Ihaka, R. G. a. R. Lexical Scope and Statistical Computing. (1996).

52 Gentleman, R. C. et al. Bioconductor: open software development for computational biology and bioinformatics. Genome Biol 5, R80, doi:10.1186/gb-2004-5-10-r80 (2004).

53 Balcells, I., Cirera, S. & Busk, P. K. Specific and sensitive quantitative RT-PCR of miRNAs with DNA primers. BMC Biotechnol 11, 70, doi:10.1186/1472-6750-11-70 (2011).

54 Busk, P. K. A tool for design of primers for microRNA-specific quantitative RT-qPCR. BMC Bioinformatics 15, 29, doi:10.1186/1471-2105-15-29 (2014).

