## Supplementary methods table for "AGO HITS-CLIP in Adipose Tissue Reveals miR-29 as a Post-Transcriptional Regulator of Leptin"

### Supplementary Tables for Methods

| KEY MATERIALS | SOURCE | CATALOG NUMBER |
|---|---|---|
| Antibodies | | |
| 2A8 Mouse mono-clonal anti-pan-Ago | Rockefeller University Antibody and Bioresource Core Facility (Note: Millipore produced 2A8, Cat#MABE56, did not produce a high efficiency immunoprecipitation) | N/A |
| Rabbit anti-mouse IgG | Jackson ImmunoResearch Labs | Cat#315005008 |
| Normal mouse IgG | Santa Cruz Biotechnology | Cat#sc-2343 |
| Commercial assays and kits | | |
| AdEasy System | Agilent | Cat#240010 |
| Dual-luciferase plasmid and kit | Promega | Cat#E1960 |
| Q5® Site-Directed Mutagenesis kit | NEB | Cat#E0554S |
| ABI high capacity cDNA synthesis kit | Applied Biosystems | Cat.# 4368814 |
| Mouse Leptin ELISA kit | Crystal Chem | Cat.# 90030 |
| Cell lines | | |
| HEK-293A cell line | Invitrogen | Cat#R70507 |
| Mouse strains | | |
| Lean C57BL/6J mice (chow-fed) | Jackson Labs | Cat#000664 |
| Obese C57BL/6J mice (HFD-fed) | Jackson Labs | Cat#380050 |
| ob$_{-/-}$ mice | Jackson Labs | Cat#000632 |
| miRNA mimics and LNAs | | |
| miR-29a miRNA mimic | Dharmacon | Cat# C-310521-07-0005 |
| Scrambled miRNA mimic | Dharmacon | Cat# CN-001000-01-05 |
| Mutated miR-29a miRNA mimic | Dharmacon | Cat# BUTMJ-000001 (seq UUGGAGCAUCUGAAAUCGGUUA) |
| miR-29a LNA | Qiagen | Cat# 339121 Y104100170-ADA |
| Scrambled ctrl LNA | Qiagen | Cat# 339126 YI00199006-ADA |
| Plasmids | | |
| Plasmid: pAdTrack | Addgene | Plasmid#16404 |
| Software and Algorithms | | |
| miRprimer2 | Busk, 2014 | https://sourceforge.net/projects/mirprimer/ |
| CTK HITS-CLIP analysis | Shah et al., 2017 | https://zhanglab.c2b2.columbia.edu/index.php/CTK_Documentation |

| | | |
|---|---|---|
| Data processing in R | Ihaka and Gentleman, 1996 | www.r-project.org |
| Bioconducter | Gentleman et al., 2004 | https://www.bioconductor.org/ |
| Miscellaneous materials | | |
| Collagenase B | Roche | Cat#11088831001 |
| Collagenase D | Roche | Cat#11088882001 |
| Dispase II | Roche | Cat#04942078001 |
| MCDB210 | Sigma | Cat# M6770-1L |
| ITS premix | BD Bio | Cat# 354352 |
| L-ascorbic acid 2-phosphate | Sigma | Cat# A8960 |
| bFGF | BD Bio | Cat# 13256-029 |
| RWT buffer | Qiagen | Cat.#1067933 |
| AdEasy bacteria BJ5183-AD-1 cells | Agilent | Cat#240010 (included in kit) |
| RQ1 DNAse | Promega | M610A |
| RNase A (20 U/ml) | Affymetrix | Discontinued |
| PowerUp SYBR Green Master Mix | Applied Biosystems | Cat.# 4367659 |
| Lipofectamine RNAiMax | Invitrogen | Cat.#13778 |

| **qPCR primers** | | |
|---|---|---|
| **mRNA qPCR** | **Oligo 1** | **Oligo 2** |
| TBP | GGGTATCTGCTGGCGGTTT | TGAAATAGTGATGCTGGGCACT |
| Leptin | TGACACCAAAACCCTCATCA | TGAAGCCCAGGAATGAAGTC |
| Pparg2 | GCATGGTGCCTTCGCTGA | TGGCATCTCTGTGTCAACCATG |
| Adiponectin | GCACTGGCAAGTTCTACTGCA | GTAGGTGAAGAGAACGGCCTTGT |
| Ap2 | ACACCGAGATTTCCTTCAAACTG | CCATCTAGGGTTATGATGCTCTTCA |
| **miRNA qPCR** | **Oligo 1** | **Oligo 2** |
| snoRNA 202 | GGCTGTACTGACTTGTAGAAAG | TCCAGTTTTTTTTTTTTTTTCATCAGA |
| miR-29a | CGCAGTAGCACCATCTGA | GGTCCAGTTTTTTTTTTTTTTTAACC |

| **Cloning PCR primers** | | |
|---|---|---|
| **Cloned gene** | **Primer 1** | **Primer 2** |
| miR-29a | (BgI11) ctaagatctATAGCTGATTAGTCAACCACC | (NotI) taagcggccgcCCACCATCACTATGTGAATAG |
| miR-16 | (XhoI) attctcgagCCTTGGAGTAAAGTAGCAGC | (HindIII) cacaagcttCAGACACAATATGTAGAGCG |
| miR-16 SDM (scrambled) | aaatattggcgTTAAGATTCTGAAATTACCTCCAGTATTGAC | acgtgttacaaAGGCACCGCTGACATTGC |

| **Luciferase assay oligonucleotides** | | |
|---|---|---|
| **Cloning gene** | **Oligo 1 (IDT)** | **Oligo 2 (IDT)** |
| Lep 1 | /5Phos/cAGTTTCGTGCTCAGCTCTGTCTGGTGCTGTGAGCc | /5Phos/tcgagGCTCACAGCACCAGACAGAGCTGAGCACGAAACTgagct |
| Lep 2 | /5Phos/cTGAGCGGGATCAGGTTTTGTGGTGCTAAGAGAc | /5Phos/tcgagTCTCTTAGCACCACAAAACCTGATCCCGCTCAgagct |

| | | |
|---|---|---|
| Mcart1 | /5Phos/cGTTTGTTTATTTTTTTAAATGGTGCTAGGGATc | /5Phos/tcgagATCCCTAGCACCATTTAAAAAAATAAACAAACgagct |
| Ptp4a1 | /5Phos/cCTTTATTAGGTTGTATATATGGTGCTAGAAGTc | /5Phos/tcgagACTTCTAGCACCATATATACAACCTAATAAAGgagct |
| Sucla2 | (/5Phos/cAGCATAGGATGTCTAGTAAATGGTGCTGGCTTc | /5Phos/tcgagAAGCCAGCACCATTTACTAGACATCCTATGCTgagct |
| Hmgcs1 | /5Phos/cCAAGTTCTCTGGATGATTTTTGGTGCTGAACAATc | /5Phos/tcgagATTGTTCAGCACCAAAAATCATCCAGAGAACTTGgagct |
| Lep 1 SDM | GCTCTGTCTGtgtaTGTGAGCCTCGAGTC | TGAGCACGAAACTGAGCT |
| Lep 2 SDM | AGGTTTTGTGtgtaTAAGAGACTCGAGTCTAGAGTCGAC | GATCCCGCTCAGAGCTCG |
| Mcart 1 SDM | TTTTTAAATGtgtaTAGGGATCTCGAGTCTAGAG | AATAAACAAACGAGCTCG |
| Ptp4a1 SDM | TGTATATATGtgtaTAGAAGTCTCGAGTCTAGAG | ACCTAATAAAGGAGCTCG |
| Hmgcs1 SDM | ATGATTTTTGtgtaTGAACAATCTCGAGTCTAGAGTC | CCAGAGAACTTGGAGCTC |
